# Hierarchical modulation of auditory prediction error signaling is independent of attention

**DOI:** 10.1101/324467

**Authors:** K. Kompus, V. Volehaugen, R. Westerhausen

## Abstract

The auditory system is tuned to detect rhythmic regularities or irregularities in the environment which can occur on different timescales, i.e. regularities in short (local) and long (global) timescale which could conflict or converge. While MMN and P3b are thought to index local and global deviance, respectively, it is not clear how these hierarchical levels interact and to what extent attention modulates this interaction. We used a hierarchical oddball paradigm with local (sequence-level) and global (block-level) violations of regularities in 5-tone sequences, in attended and unattended conditions. Amplitude of negativity in the N2 timeframe and positivity in the P3b timeframe elicited by the final tone in the sequence were analyzed in a 2*2*2 factorial model (local status, global status, attention condition). We found a significant interaction between the local and global status of the final tone on the N2 amplitude (p<.001, η_p_^2^= .55), while there was no significant three-way interaction with attention (p > .05), together demonstrating that lower-level prediction error is modulated by detection of higher-order regularity but expressed independently of attention. Regarding P3b amplitude, we found significant main effect of global status (p<.001, η_p_^2^= .42), and an interaction between global status and attention (p < .001, η_p_^2^= .70). Thus, higher-level prediction error, indexed by P3b, is sensitive to global regularity violations if the auditory stream is attended. The results demonstrate the capacity of our auditory perception to rapidly resolve conflicts between different levels of predictive hierarchy as indexed by MMN modulation, while P3b represents a different, attention-dependent system.

## Introduction

Our perception relies on prediction to facilitate the decoding of the sensory information. The predictive coding theories of perception suggest that the brain tries to minimize the surprise or prediction error, and continuously uses the unpredicted portion of the sensory input to adjust the predictive models (1). A crucial component of the predictive coding is the hierarchical organization of perceptual systems, with higher levels which represent slower-changing regularities modulating the processing of lower-level predictive units which integrate over shorter time (2). Such nested hierarchical system are crucial in human speech processing, where the probability of a sound depends on its immediate local environment such as the syllable structure, whereas word and sentence rules in the given language give wider context which would need to be taken into account when predicting the subsequent sound (3). How such hierarchically nested rules are extracted from the auditory stream, how they interact with each other and other systems such as attention and long-term memory are central to our understanding of auditory perception.

Unpredictable deviations from predicted pattern in auditory input are detected and given special processing in the auditory analysis of the incoming sounds. This can be detected using event-related potentials, where mismatch negativity (MMN), a negativity best visible in the difference wave between a predicted and an unpredicted stimulus, arises 150-250 ms after the onset of the deviant (4). While neuronal adaptation contributes to the MMN generation, the MMN cannot be fully explained by adaptation as it can be elicited by a deviant which is physically identical to the standard but violates an otherwise established rule (5, 6). MMN has been suggested to reflect the prediction error which results when a perceptual predictive model meets sensory input (7). The portion of the input which is compatible with the model will be attenuated, whereas the unpredicted portion will be propagated further and used to update the predictive model. This would explain why physically more different stimuli, as well as less probable stimuli lead to larger MMN: there capacity of the model to explain the incoming information is lower, and consequently larger error variance will be propagated (7, 8).

Most MMN studies have examined the predictability of a stimulus with regard to its immediate environment: the immediately preceding sounds, presented with relatively short interstimulus interval (<1000 ms), where the predictive model would predict continuation of the pattern established by the immediately preceding stimuli (9). However, in addition to such “local” rules which govern whether the stimulus is predictable from a short train of preceding stimuli, there could be more general rules about the predictability of the current stimulus in a wider context, which could agree with or diverge from the local rules. For example, a stimulus which is locally unpredictable, deviating from several preceding identical stimuli, could be following a rule establishing its regular occurrence over a longer time period. Such “global” rule would take into account the probability of the stimulus in a more complex situation. Paradigms which have examined such “local”/”global” rules have used a set-up where two stimuli, A and B, are presented in regularly structured sequences (e.g., AAAAB), so that the model could build up an expectancy about a physical deviant, which could be rarely violated by a physical standard (e.g., by a sequence AAAAA) (10). Such pattern extraction-MMN (11) has elicited considerable interest as it addresses the question how the auditory system can efficiently represent the often complicated and conflicting rules governing the probability of stimuli.

It has been suggested that simple tone *vs.* complex pattern deviations, corresponding to violating the local and the global rule, respectively, are processed by different neural systems, with simple-feature deviance-detection occurring at earlier levels of auditory processing and increasingly complex rule deviations detected on higher levels (12). An important question, however, is whether the higher levels can modify the processing in the earlier levels, as suggested by the hierarchical coding model (1): when the global rule has been learned, can it modify the detection of the deviance due to the local rule, and suppress the MMN for a stimulus which differs from its immediate environment but is predictable according to the global rule?

The MMN generation to an unpredictable stimulus also appears to be connected to the distribution of attentional resources: an unexpected deviance which the subjects can detect typically elicits not only the MMN but also ERPs associated to attention, such as the P3a component which has been linked to general novelty-processing and attention-switching mechanisms (13). However, the link between MMN and attentional processes does not appear to be directly causal: the P3a is not an obligatory sequel to MMN, but instead appears to depend on the level of MMN or be linked to N1 instead (14–16). Another aspect of the MMN-attention relationship is the question whether attentional resources are required for deviance detection to unfold in the first place. While MMN is often referred to as attention-independent, relying on the observation that a robust MMN is elicited when the subjects are attending visual stimuli (5), it has been proposed that attention can reorganize the auditory stream before it is passed on to the processes which lead to MMN generation (17). One interesting situation where such attention-dependent reorganization could have a substantial effect is the detection of regularities which unfold over a longer time period: seconds or even minutes.

Crucially, however, global-rule formation has been suggested to be attention-dependent. An influential series of studies (10, 18–20) has used a local/global paradigm where five-tone sequences (AAAAA or AAAAB) are presented within a longer block of sequences, where each sequence can be either typical to the block (i.e., majority of the sequences are the same type) or differ from the rule imposed by the majority within the block. In this situation two hierarchically distinct predictions can be made: local predictions about the fifth tone in the
sequence, and global predictions about the probability of the fifth tone while taking into account the probability of the five-tone sequence in the context of the current block. These studies have suggested that there is interaction between the global deviance processing and attention: while the local rule representation is done automatically and independently of attention, detecting the deviance from the global rules is performed by a system which is dependent on the ability to dedicate conscious attention to the auditory input. With this paradigm, the violation of the global rules (the five-tone sequence differs from a rule of the group) has been noticed to elicit P3b component, which is not elicited when the subjects are not actively attending the stimuli. This has led to the suggestion that the global rule representation is performed in different neural structures, which are indexed by the P3b generation, and the operation of these structures is reliant on the availability of the global workspace. This attention-dependent higher-order rule can then, when it has been formed, regulate the “local” effect which is elicited in the MMN timeframe (21). Such dual-process view with attention-dependent context processing, however, conflicts with the studies that demonstrate that violation of an established pattern of sounds which unfolds over longer time interval elicits MMN even in the absence of conscious awareness. For example, (22) found that a pattern-violation MMN was generated by a deviation from a four-tone pattern even when the subjects did not consciously notice the pattern; see also (23).

An important examination of the relationship between the supposedly attention-dependent global rule representation and automatic local rule representation is whether the modulation of the local by the global rule (i.e., the interaction in the MMN timeframe) is present even when the subjects are not paying attention to the stimuli. This has not been systematically examined in the studies which have proposed the different neural underpinnings of the local and global rule representation (18–21). While several studies showed that attention did not interact with the local deviance effect, but did interact with the global deviance effect (10, 18, 19), this does not, however, address the question whether the global rule is formed and affects the MMN processes in the absence of attention. The effect of interest is the interaction between local and global factors in the MMN time window, as this demonstrates whether the hierarchically higher global rule can influence the lower-level local rule. Consequently, to examine whether attention affects the global rule representation, the three-way interaction between attention, local and global factors in the MMN time window needs to be tested.

In this study we used the local/global paradigm under attended and unattended condition to examine whether the interaction between the local and global predictions is modulated by whether or not the stimuli are attended. In case the interaction between the local and global rules in the MMN timeframe is affected by the attention, this suggests that the extraction of a global regularity from an auditory stream is indeed attention-dependent. By contrast, in case the attention only affects the P3b elicitation but not the local-global interaction within the MMN timeframe, it suggests that the global rule extraction is performed without attention, and the P3b represents a different process which is attention-dependent, but independent from the global rule representation in the auditory stimulus processing.

In addition to N2 and P3b, we also examined the effect of the local and global rule violation under attended and unattended conditions on the P3a, a component with a frontocentral maximum following the negativity in the MMN timeframe, associated with the attention capture by the deviant stimulus. The P3a has been suggested to index the attentional capture by the deviant stimulus (13), and appears to be generated when the MMN reaches a certain threshold value. If the P3a demonstrates interaction of the local and global predictions, this would show that the attentional capture is either directly coupled to the magnitude of the MMN, or sensitive to the same contextual modulation as the MMN. By contrast, a lack of interaction would support the view that P3a generation is not directly coupled to the process that creates MMN. Further, by looking at the interaction with attention we can test whether the attention to the stimuli gates the attentional shift, or whether it is independent of the attention.

## Methods

### Participants

The participants were 20 healthy young adults (10 female) recruited among the community of the University of Bergen. Three subjects’ data was discarded due to excess movement artefacts during EEG session, the analyses are based on 17 subjects (7 female). The mean age of the 17 subjects was 25.4 years (range 21.1-34.8, sd 3.6). Subjects had no history of psychiatric or neurological disease (assessed by self-report) and had auditory thresholds <25 dB in both ears for frequencies 250, 500, 1000, 2000 and 3000 Hz as assessed using the Hughson-Westlake audiometric test (Oscilla USB-300, Inmedico, Lystrup, Denmark). The subjects were asked to refrain from nicotine and caffeine for at least 5 hours before the recording. The participants were informed about the experimental procedures and signed an informed consent form. The study was approved by the regional ethics committee (REK-VEST)

### Paradigm

The “Local-Global” MMN paradigm consisted of sequences of five tones. The tones were harmonic tones composed of three sinusoidal partials (tone A: 500, 1000, 1500 Hz; tone B: 550, 1100, 1600 Hz) with 50 ms duration (including 7 ms rise and fall time). Interval between tones in a sequence was 100 ms, with total sequence length 650 ms. The sequences were either XXXXX (five identical tones, AAAAA and BBBBB) or XXXXY (fifth tone different, AAAAB and BBBBA). The final tone in the sequence could thus be either local standard or local deviant.

The sequences were presented in blocks, with inter-sequence interval 700-900 ms (random variation with step 50 ms, mean 800 ms). Half of the blocks had 80% XXXXX sequences and 20% XXXXY sequences, in the other half the frequency was reversed (80 % XXXXY sequences, 20 % XXXXX sequences). The sequences could thus be global standards (the dominant sequences) or global deviants (the rare sequences). This leads to a factorial design to probe the final tone of the sequence (global status: deviant/standard, local status: deviant/standard).

Each block began with 25 repetitions of the sequence which was the global standard for that block, serving as a model-building phase, following which 150 sequences were presented (in 80/20 ratio). The block length was 4 minutes 22 seconds. Inter-block interval was 3 seconds. In total, 8 blocks were presented: four with XXXXX as global standard and four with XXXXY as global standard.

Two attentional conditions were used: attended and unattended stimulation. In the attended condition the subjects were asked to monitor the tones, compliance was checked by asking the subjects to report on the sound characteristics after the recording using a 5-item questionnaire. All subjects reported detecting some of the regularities present in the sound streams, with the average score 4.1 out of 5 possible (only three subjects scoring 2 or 3, the others 4 or 5). In the unattended condition the subjects were asked to perform a visual working memory task. In the visual n-back task abstract visual objects (Fribbles, TarrLab, http://www.tarrlab.org.) were presented, asynchronously with the auditory stimuli, and the subjects asked to press a button in case the object was the same as 2 objects previously. Compliance was checked by examining the response profile of the visual n-back task. All subjects gave responses to the visual task. The mean accuracy (proportion of hits and correct rejections) was 0.87 (range 0.77-0.95, sd 0.06); the mean sensitivity index d’ was 1.65 (range 0.82-2.74, sd 0.62).

### EEG recording and analysis

The data were recorded in an electromagnetically shielded EEG recording chamber using 12 Ag/AgCl electrodes (F3, Fz, F4, FCz, C3, Cz, C4, TP9, TP10, P3, Pz, P4) placed according to the International 10-20 system using the EasyCap electrode cap (Falk Minow Services, Breitenbrunn, Germany) and Abralyt 2000 electrode gel. Interelectrode impedance was kept under 10 kΩ. The reference electrode was placed at nosetip, the ground at FT10. Four electrodes were used for monitoring eye movements, two placed above and below the right eye and at the outer canthi of the eyes. The EEG data were recorded using BrainVision Recorder 1.0 (Brain Products, Munich, Germany). The sampling rate was 500 Hz, filter band 0-100.

After recording, the data were offline filtered using a zero-phase Butterworth IIR filter with high-pass threshold 0.01 Hz (slope 12 dB/oct) and low-pass threshold 30 Hz (slope 12 dB/oct). The data were downsampled to 250 Hz. Eye movements were removed using Gratton-Coles algorithm implemented in the BrainVision Analyzer. The data was epoched into segments relative to the onset of first tone in the 5-tone sequence. Epochs spanned from - 100 to 1348 ms after the onset of the first tone, covering the entire 5-tone sequence. Automatic artefact detection was used, with epochs where the amplitude exceeded +-75 μV were discarded. The final tone (which could deviate from the local or global rule) onset was at 650 ms.

To reduce the effect of contingent negative variation between attended and unattended conditions, which could have affected the waveform from the beginning of the first tone, the epochs were corrected to the mean of the 50-ms baseline before the onset of the final tone. Relative to the onset of the final tone, the N2 was quantified as the most negative value (averaging over +/- 20 ms around the peak) in the time window 50-250 ms in the electrode FCz; the mastoid positivity was examined as the most positive value in the same time window at the electrodes TP9 and TP10. We tested the N2 peak in the raw average waveform, for comparability with the previous studies using the local-global paradigm as used here. However, to reduce the contribution of other, attention-dependent potentials in the same timeframe, we also calculated the difference waves between the standard and global sequences and examined the MMN as a negativity evident in the difference wave between the 3 types of deviant stimuli (local-only, global-only, local and global) and the standard stimulus (local and global standard). The three difference waves in both attended and unattended conditions were calculated, and for each subject the most negative value in the time window 50-250 ms was extracted (mean around +/- 20 ms surrounding the peak).

The P3a was quantified as the most positive value following N2 up to 400 ms post-deviant onset at the electrode FCz. Finally, the P3b was quantified as the area under the curve in the interval 300-500 ms at the electrode Pz.

### Statistical analysis

Extracted peak value analysis was performed on IBM SPSS Statistics version 25. The data were analyzed using a 2*2*2 repeated-measures general linear model, with factors Local status (standard/deviant), Global status (standard/deviant) and Attention (attended/unattended). The effect sizes were calculated as partial eta squares (η_p_^2^), for post-hoc paired comparison the Cohen’s d were calculated. The effects of interest are the two-way interaction between Local and Global factors, and the tree-way interaction between Local, Global and Attention for the peak values in the time window 50-250 ms (N2) as well as P3a time window, testing the contextual modulation of local deviance detection by N2 and P3a, respectively, and testing whether the contextual modulation of these peaks is affected by attention. In addition, in the P3b time window the effect of interest is the main effect of Global and interaction between Global and Attention, testing the sensitivity to the global rule as well as the attention-dependence of global rule representation by the P3b.

In addition to the windowed analysis in pre-selected time intervals, we also estimated the GLM as specified above at each timepoint after the onset of the final tone to examine the time course of the attentional effects on the local and global status and their interaction on the ERPs. For significance testing we used the PALM package (Winkler et al., Neuroimage 2014) with the threshold-free cluster enhancement procedure, performing 8000 permutations; the p-values were corrected using familywise error correction.

## Results

### N2 and MMN

The waveforms for N2 as well as difference waveforms isolating the MMN component are depicted on Figure 1, row A and B. As can be seen in the difference waveforms (Fig.1, B), a clear MMN was present in all conditions; testing the peak amplitude at FCz electrode against zero indicated that the peak value was significantly below zero in all six conditions (all p<.005). We then examined the effect of the different experimental conditions on the N2 peak (Fig.1, A) using the factorial general linear model as described above. In FCz, there was a main effect of Local status (F(1,16)=22.4; p<.001, η_p_^2^= .85, Global status ((F(1,16)=87.1; p<.001, η_p_^2^= .58) and Local*Global interaction (F(1,16)=19.6; p<.001, η_p_^2^= .55). There were no significant main effects or interactions involving the Attention factor. Post-hoc pairwise comparisons, collapsing over the Attention factor, showed that the standard *vs* deviant effect for the Local factor depended on the level of the Global factor: for items that were also global standards, the local standard-deviant difference was smaller (sta: -.93, dev: -1.45; t (16) =-1.6, p=.12, d=.36) than for items that were also global deviants (sta: -1.9, dev:-4.5; t(16)=-5.5, p<.001, d=1.5).

**Figure. 1.**
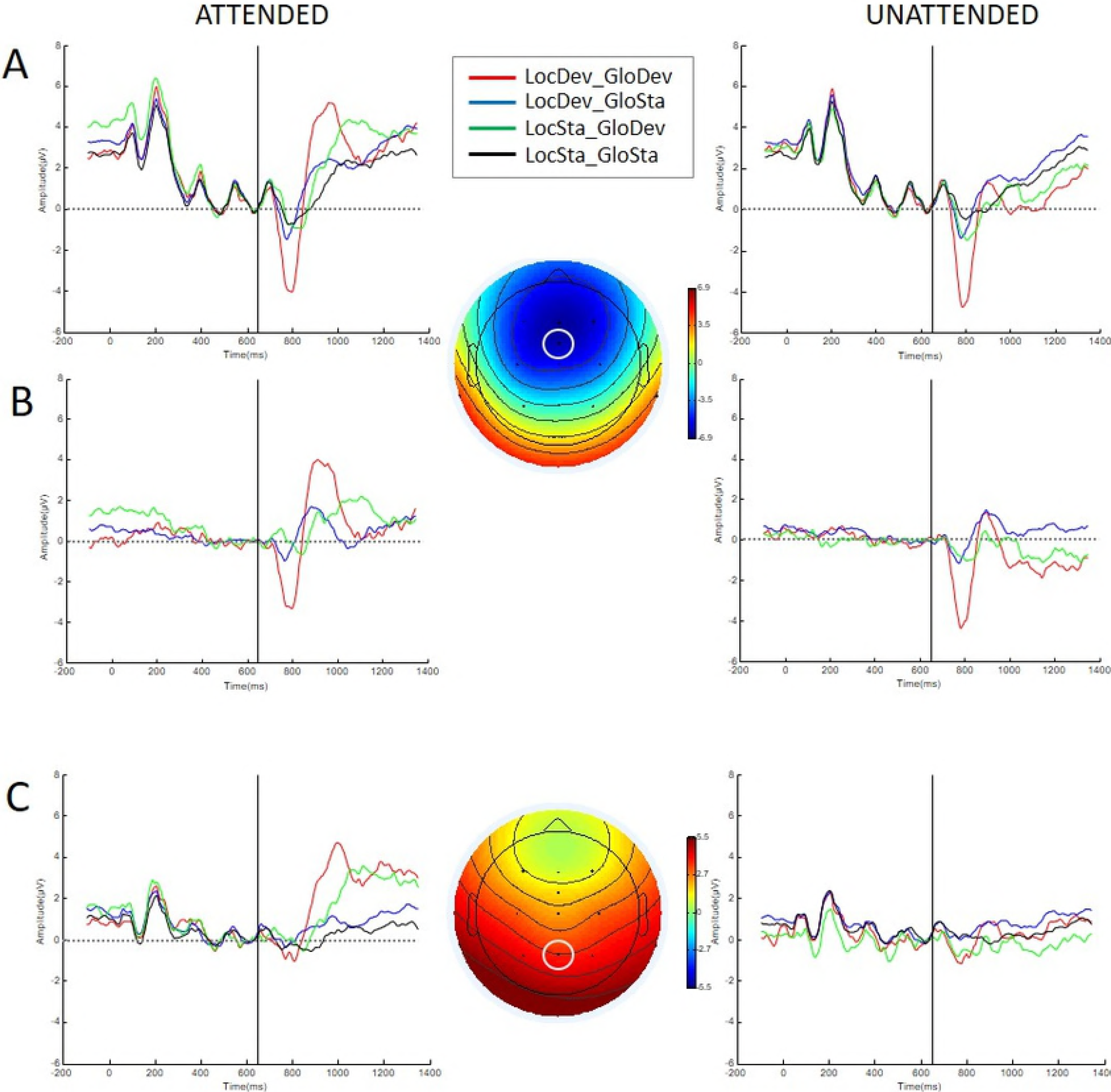
A: The waveforms from FCz depicting the N2 and the P3a for attended (left) and unattended (right) condition. B: The difference waveforms from FCz for the three deviant condition relative to the local and global standard condition, isolating the MMN component. All MMN peaks are significantly below zero (see text). The GLM with factors Attention and Type in FCz showed significant main effect of Type (F(2,32)=33.2, p<.001), with no significant main or interaction effects of Attention. Pairwise comparison between the three types showed significant difference between local-global and the other two types. C: The waveforms from Pz depicting the P3b for attended (left) and unattended (right) condition.

For the mastoid electrodes, additionally the Laterality factor was included in the GLM to assess whether the effects differ between left and right side. As there was a three-way interaction between Laterality, Local and Global (F(1,16)=14.0, p=.002, η_p_^2^= .47), we performed separate GLMs in each of the electrode sites. In the left mastoid, there was a significant main effect of Local status (F(1,16)=18.9, p=.001, η_p_^2^= .54), Globl status (F(1,16)=58.5, p<.001, η_p_^2^= .79) and Local*Global interaction (F(1,16)=29.3, p<.001, η_p_^2^= .65). Same pattern was found in the right mastoid: there was a significant main effect of Local status (F(1,16)=10.5, p=.005 η_p_^2^= .40), Global status (F(1,16)=13.8, p=.002 η_p_^2^= .46) and Local*Global interaction (F(1,16)=7.7, p=.014 η_p_^2^= .33). There were no significant main effects or interactions involving the Attention factor in either of the mastoid electrodes. The effect sizes suggest that the three-way interaction involving the electrode factor was due to difference in effect sizes, but not due to differences in the overall pattern of associations. Post-hoc pairwise comparisons, collapsing over Attention factor, showed same pattern as in FCz: the standard vs deviant effect for the Local factor depended on the level of the Global factor. In the left mastoid, for items that were also global standards, the local standard-deviant difference was smaller (sta: 1.1, dev 1.3, t=1.11, p=.28, Cohen’s d: 0.28) than for the items that were also global deviants (sta: 1.8, dev: 3.34, t=5.5, p<.001, Cohen’s d: 1.3). In the right mastoid, similar pattern was seen: for items that were also global standards, the local standard-deviant difference was smaller (sta: .66, dev: 1.03, t=2.09, p=.05, Cohen’s d: .53) than for the items that were also global deviants (sta: 1.3, dev 2.28, t=3.4, p=.004, Cohen’s d: .91).

### P3a

The waveform demonstrating P3a is depicted on Figure 1 (A). There was a main effect of Local status (F(1,16)=6.6, p=.02, η_p_^2^= .29) and Global status (F(1,16)= 14.5, p=.002 η_p_^2^= .48), with deviant stimuli eliciting larger amplitude than standard stimuli, and Attention (F(1,16)=10.9, p=.005 η_p_^2^= .41), with attended stimuli eliciting larger amplitude than unattended stimuli. The Local*Global interaction as well as Local*Global*Attention interaction were not significant. However, there was a significant interaction between Attention*Global (F(1,16)=16.7, p=.001 η_p_^2^= .51). Post-hoc pairwise comparisons, collapsing over the Local factor showed that the global-deviants and global-standards differed only in the attended condition (5.58 vs 2.7, t=4.6, p<.001, Cohen’s d: 1.6); while there was no significant difference in the unattended condition (1.99 vs 1.83, t=.42, p=.68, Cohen’s d: 0.11).

### P3b

The waveform showing P3b in the electrode Pz is depicted on Fig.1, C. There was a significant main effect of Global status (F(1,16)=11.6, p=.004; η_p_^2^= .42), Attention (F(1,16)=13.8, p=.002; η_p_^2^= .46), and a significant interaction between Global status and Attention (F(1,16)=38.0, p<.001; η_p_^2^= .70). Post-hoc pairwise comparisons, collapsing over the Local factor showed that the global-deviants compared to global-standards were more positive only during attended condition (3.2 vs .68, t=5.4, p<.001, Cohen’s d: 2.2), whereas during unattended condition the deviants were slightly more negative than standards (-.07 vs .52, t=-2.2, p=.04, Cohen’s d: -0.58). Other effects were not significant.

### Timepoint-by-timepoint analysis

The results of the permutation tests are shown on Figure 2. They indicate that the Local*Global interaction effect demonstrated in the peak analysis above extends across the frontal and central electrodes, stronger on the midline and right, whereas the Attention*Global interaction shown for P3b encompasses not only the parietal but also central electrodes and lasts from approximately 300 ms post-deviant onset to the end of the epoch. The timepoint-by-timepoint analysis also confirms that the three-way interaction between local and global status and attention is not significant anywhere, at most a weak trend can be seen in a small number of timepoints in electrodes F3 and C3. Also, attention does not interact with the Local factor anywhere, supporting the notion that MMN generation is not modulated by attention, at least in situations where attention does not lead to substantial reorganization of the auditory stream. Of note is a discrepancy between Local and Global factors in the P3a: while the Local deviants lead to a significant effect in the P3a range starting quite early (~200 ms post-deviant and centrally located, suggesting that it is the early, attention-independent phase of the P3a), by contrast the effect did not survive the familywise error correction for the Global factor. The later, 300-ms onset effect in frontocentral and central electrodes visible in Attention*Global plot could be related to the later phase of the P3a. Finally, it is noticeable that both Local and Global factors have a significant effect in the bilateral mastoid electrodes after ~250 ms post-deviant onset, and the effect extends longer in the Global than Local. This could indicate the auditory cortex-level model readjustment.

**Figure. 2.**
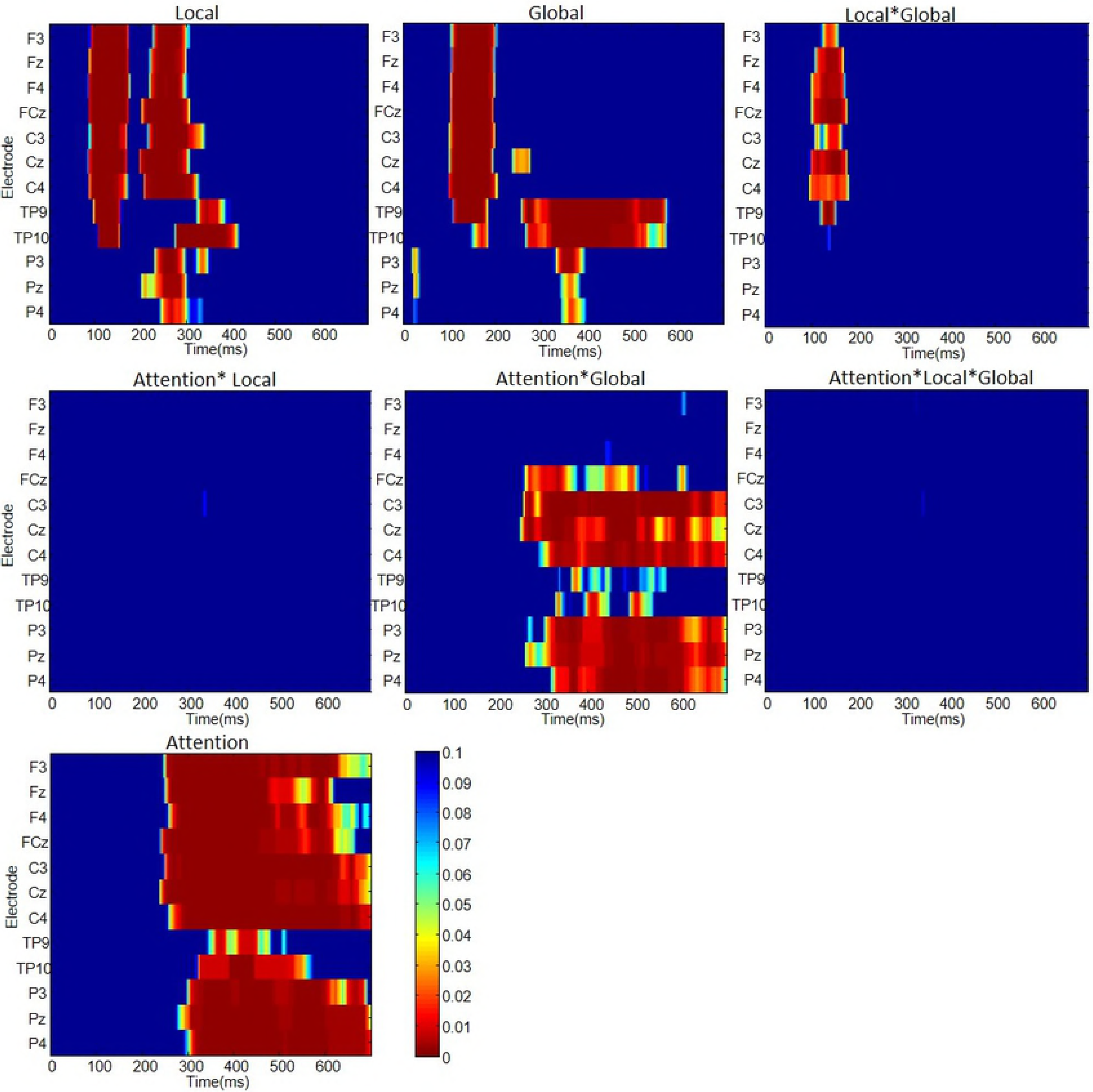
The timepoint-by-timepoint permutation analysis, showing p-values (family-wise error corrected) for the effects of the factors Local, Global, Attention and their interactions. No timepoints were significant for Attention*Local and Attention*Local*Global analysis.

## Discussion

In this experiment we exposed subjects to a hierarchical auditory structure where the frequency of a tone was predicted by two independent rules in a factorial design: a local rule in relation to immediate environment and a global rule applying over longer timeframe. The local rule violation as well as the global rule violation elicited a negativity in the N2 timeframe, having the characteristics of a classical MMN response (frontal negativity and simultaneous mastoid positivity). As has been observed in earlier studies (10, 18, 20, 21), the global rule violation elicited a P3b wave only in the attended condition, confirming that the attention manipulation was successful. However, despite the absence of P3b for unattended global deviants, there was a significant interaction between the local and the global rule in the MMN timeframe which did not interact with the attention factor, indicating that even though the global rule violations were not registered by the P3b-generating system, they influence the MMN. The results demonstrate that the processes indexed by the negativity in the MMN timeframe are sensitive to both the local as well as the global status of a sound. If the predictions made by observing the local and the global level transitional probability agree, the amplitude of the negativity is either large (unpredictable on both levels) or small (predictable on both levels). However, if the transitional probabilities predicted by the two levels conflict, the resulting probability is intermediate, as if the two levels would cancel each other out. This interaction between the local and global predictions indicates that the two are carried out by different neural correlates which interact within the MMN timeframe.

It is also evident that the interaction of these local and global predictors is not sensitive to attention, as there was no significant interaction effect with attention. This further suggests that the formation of the global rule does not require conscious attention, in contrast to claims by earlier studies (10, 18, 20). We demonstrated that the ERPs in the MMN range are sensitive to the modulation by the global probability of a five-tone sequence in its block, even while the subjects were attending a visual working memory task. This indicates that the representation of auditory environment in the order of several seconds and consisting of more complex patterns is carried out by the brain without needing the conscious attentional resources. The pattern of significant main effects of local and global as well as interaction between local and global factors seen at the frontocentral electrode was replicated in the mastoids, indicating that the observed pattern was not due to contamination from any other attentional effects which are typically expressed in the frontocentral, but not in the mastoid locations (24).

An alternative interpretation of the results is the differences in the transitional probability of the tones across different conditions. The local deviant consisted of a physically different tone relative to the four preceding tones (XXXXY sequence). The interaction effect indicates that the ERP elicited by the final Y-tone had a different amplitude in the XXXXY block, where Y-tones made up 16 % of all the tones, compared to the XXXXX block, where Y-tones made up 4 % of all the tones. Thus the difference in the amplitude would be consistent with the observation that the standard *vs*. deviant difference increases with reduced overall probability of the deviant tone (5). By this interpretation, the effects would not be due to a global rule interacting with a local rule, but could represent factors such as stronger stimulus-specific adaptation in the XXXXX blocks. However, such explanation does not agree with the effect of Global status on the negativity in the MMN timeframe in the XXXXX sequences. While locally standard, the N2 amplitude to the final x-tone differed depending on whether the sequence was embedded in an XXXXX-block or XXXXY-block, an effect also seen when examining the difference waveforms isolating the MMN component (Fig.1). The overall probability of the x-tone in these different blocks was 96 % and 84 % respectively, which means it would have been a local standard in both conditions. Thus, the modulation of the negativity by the sequence-final X-tone could not be due to the local transitional probability relative to the previous tone but reflected the global rule of the probability of the XXXXX-sequence relative to the block. This finding agrees with the literature demonstrating complex and long-timewindow regularity representation in the auditory cortex, as indexed by MMN (11, 23).

We additionally characterized the MMN wave generated by the deviants relative to the standard sequence. The difference waves relative to sequence which was both a local and a global standard (Fig.1, B) show the expected significant negativity in all three deviant conditions.

The parallel representation of the local and global rule is consistent with the studies on multi-feature MMN, which have looked at the effect of violating predictions regarding different physical features of a sound independently (simple deviation) or in conjunction (abstract deviation). As demonstrated by Takegata et al. (25), representation of the simple and abstract rules was carried out in parallel, and the simultaneous deviation from both of these generated a MMN waveform which was explained by a model of combined effect of the two types of rules. In the present case, the effect was interactive rather than additive, but the parallel representation of two rules in predicting the same feature is in agreement with this model. The present results add the perspective that the more higher-order rule may consist of auditory Gestalt representation with relatively long time-window, as the representation of the global rule here needed to maintain a representation of at least the previous 5300 ms to hold two repetitions of the standard sequences (considered to be the minimum standard-building requirement) and react to the deviation in the final tone of the subsequent sequence.

In contrast to the N2, the results in the P3a and P3b showed that their representation of violation of predictions was modified by attention. There was a slight divergence in their response. P3b was not sensitive to local status of the sound. This has in earlier literature led to suggestion that P3b is uniquely representing the higher, block-level status of incoming information, which is dependent on attention, and is unable to operate when the attentional resources are removed (10, 18, 20). The current results demonstrate that this is not the full picture. P3b is indeed sensitive to interaction between global status and attention: the global deviants and standards differed only in the auditory attention condition, and did not differ in the unattended condition when the subjects were performing a visual n-back task as a distractor. However, as discussed above, the modulation of the N2 by the global rule was clearly present in the attended as well as unattended condition. This indicates that the processes leading to negativity in the MMN timeframe could track the global status of the sound even under conditions where the P3b was not capable of representing it. Therefore, the data does not support the view that global rule representation is uniquely performed by a system where violation generates P3b. Instead, the P3b appears to index conscious, attentional processing of the attended sequences. This is consistent with the theories interpreting P3b as an index of detecting events which are salient or important relative to the currently maintained goal state (26, 27).

The P3a, unlike P3b, was sensitive to the local status of the tone, similarly to N2. This is in agreement with findings of P3a following the violations which elicit the MMN. The P3a, linked to frontal lobe as well as medial temporal lobe, is frequently associated to novelty processing. In the context of auditory oddball studies it has been associated to orienting attention toward the deviant sound, or evaluating the contextual novelty of sounds (13). However, the effect was not modulated by the global rule, corroborating the suggestions that P3a generation is not directly dependent on the amplitude of MMN, instead it depends on processes happening prior to MMN generation. Also, there was no significant interaction between Local and Attention factors: both attended and unattended local deviants elicited P3a with similar amplitude. However, similarly to P3b, there was an interaction between global status and attention: the effect of the global status on the amplitude change during deviant compared to standard was larger under the attention condition than visual distractor condition. As indicated by the timepoint-by-timepoint permutation analysis, this may be due to two effects overlapping in the same time-window in the FCz electrode: an earlier Local effect which starts at ~200 ms post-deviant onset and extends from frontal to central electrodes and a later Attention*Global effect which begins ~250-300 ms post-deviant onset and is confined to frontocentral and central locations. Considering the distribution, it seems plausible that the early, post-200 ms component of the P3a, which is attention-independent and with more frontal location (13) is distinct from the later-onset component which appears to behave more similarly to P3b.

The effect of attention on prediction error processing is debated. Kok et al. (28) and Garrido et al. (29) explored two possibilities: attention and unpredictability each lead to increased processing, but there is no interaction between them; vs an interactive model where attention “flips” the preference so that under attention, predicted stimuli receive more processing, whereas in unattended condition, unpredicted stimuli are processed more. Garrido et al. (29) showed that for auditory stimuli the model where attention and prediction give independent boosting effects is preferred over interaction-model. The present study similarly suggests that the interaction of attention and prediction on early sensory processing shown by Kok et al. (28) is not robustly found in all experimental situations, and may thus reflect effects specific to visual processing. The present results are in agreement with Garrido et al. (29), in demonstrating independence of attention and prediction in the N2 time range. There was a main effect of Attention across all EEG channels from ~250 ms after the onset of the final stimulus, however, the local unpredictability did not show interaction with this effect, and led to effects around 150 ms. While the global predictions and attention interaction was seen from 250 ms in central and parietal electrodes, the effect was characterized by increased processing of unpredicted sequences compared to predicted sequences, opposite to the prediction made by (28), while typical for oddball studies on P300 ERP. While attention and prediction interaction at 190 ms has been reported by (30) in an MEG study, that study had low number of deviant trials and found the interaction effect in the right temporal sensors following the main mismatch negativity response, thus possibly rather reflecting processes related to the P3a.

## Conclusion

An important finding of this paper is demonstrating the independence of the P3b and the global rule representation. While the global deviants compared to global standards elicited P3b wave only in the attended condition, similarly to previous reports, the modulation of the N2 by the global rule was clearly present in the attended as well as unattended condition. This was visible both when considering the N2 peak as a simple or difference waveform, revealing MMN and removing any other components which may be expressed in the same time window. The data show that in the local-global paradigm the P3b elicitation marks the presence of deviant sound sequences when asked to attend the sounds, but the global status affects the sound processing already earlier, in the N2 window where the MMN is typically expressed. While the N2 window demonstrates interaction between the local and global rule representation, this is independent of attention. This adds to the body of literature showing that the MMN process in the N2 time window is robust and does not show sensitivity to central executive load.

